# easier: interpretable predictions of antitumor immune response from bulk RNA-seq data

**DOI:** 10.1101/2021.11.26.470099

**Authors:** Óscar Lapuente-Santana, Federico Marini, Arsenij Ustjanzew, Francesca Finotello, Federica Eduati

**Affiliations:** Department of Biomedical Engineering, Eindhoven University of Technology, 5612 AZ Eindhoven, The Netherlands; Institute of Medical Biostatistics, Epidemiology and Informatics (IMBEI), University Medical Center Mainz, D-55131 Mainz, Germany; Institute of Molecular Biology, University of Innsbruck, 6020 Innsbruck, Austria; Digital Science Center (DiSC), University of Innsbruck, 6020 Innsbruck, Austria; Institute for Complex Molecular Systems, Eindhoven University of Technology, 5612 AZ Eindhoven, The Netherlands

## Abstract

Immunotherapy with immune checkpoint blockers (ICB) is associated with striking clinical success, but only in a small fraction of patients. Thus, we need computational biomarker-based methods that can anticipate which patients will respond to treatment. Current established biomarkers are imperfect due to their incomplete view of the tumor and its microenvironment. We have recently presented a novel approach that integrates transcriptomics data with biological knowledge to study tumors at a more holistic level. Validated in four different solid cancers, our approach outperformed the state-of-the-art methods to predict response to ICB. Here, we introduce **e**stim**at**e **s**ystems **i**mmun**e r**esponse (*easier*), an R/Bioconductor package that applies our approach to quantify biomarkers and assess patients’ likelihood to respond to immunotherapy, providing just the patients’ baseline bulk-tumor RNA-sequencing (RNA-seq) data as input.

## 1 Introduction

Immunotherapy with immune checkpoint blockers (ICB) exploits the existing patient’s immunity to attack tumor cells (Waldman et al., 2020) and demonstrated striking clinical efficacy but just in a minority of patients (Sharma et al., 2017). Thus, predicting patients’ response to ICB based on biomarkers is a pressing need in immuno-oncology (Tang et al., 2018).

Tumors are multicellular systems governed by a complexity of regulatory interactions within cancer cells, but also with the surrounding tumor microenvironment (TME). Approaches that take a comprehensive view of the TME are strongly required to understand immune response and, to identify interpretable, mechanistic biomarkers (Lapuente-Santana and Eduati, 2020; Du and Elemento, 2015). We recently developed a novel approach that uses machine learning to combine transcriptomics data with prior information on TME, to derive system-based signatures that provide a mechanistic understanding of the tumor and effectively predict patients’ response to ICB (Lapuente-Santana et al., 2021).

This pipeline (Supplementary Fig. S1) is now implemented in *easier* (**e**stim**a**te **s**ystems **i**mmun**e r**esponse), an R/Bioconductor package. The package requires as input bulk RNA-sequencing (RNA-seq) data, and allows to derive the quantitative signatures of the TME and to use pre-trained cancer type-specific machine learning models to predict patients’ immune responses. *easier* also allows to integrate information on tumor mutational burden (TMB). The new *easier* R package provides a likelihood score of antitumor immune response and access to the derived biomarkers.

## 2 The easier package

*easier* integrates bulk RNA-seq data with different types of prior knowledge to extract mechanistic quantitative descriptors of the TME (called *features* hereafter). Leveraging the large datasets from The Cancer Genome Atlas (Cancer Genome Atlas Research Network et al., 2013), we employed multi-task linear regression to learn how these features could predict different hallmarks of antitumor immune response and to derive cancer type-specific models based on interpretable biomarkers (Lapuente-Santana et al., 2021). *easier* uses these trained models to identify system biomarkers of immune response and to predict patients’ likelihood of response to ICB using patients’ baseline bulk RNA-seq data as input.

Input data can be used to compute five quantitative descriptors of the TME: immune cell fractions (quanTIseq; Finotello et al., 2019), pathway activity (PROGENy; Schubert et al., 2018), transcription factor activity (DoRothEA; Garcia-Alonso et al., 2019), ligand-receptor and cell-cell interaction scores (Lapuente-Santana et al., 2021).

These features are used as input for our cancer type-specific models to obtain patients’ predicted immune response (Supplementary Fig. S1). *easier* provides a likelihood of immune response and a corresponding confidence interval for each patient (Supplementary Fig. S2). When the information about patients’ response to therapy is available, *easier* additionally allows evaluating the predictions against the ground truth. *easier* offers the possibility to further integrate information on TMB to derive an integrated score to account for the complementary role of immune response and tumor foreignness for successful ICB therapy (Fig. 1A; Supplementary Fig. S3).

**Fig. 1.**
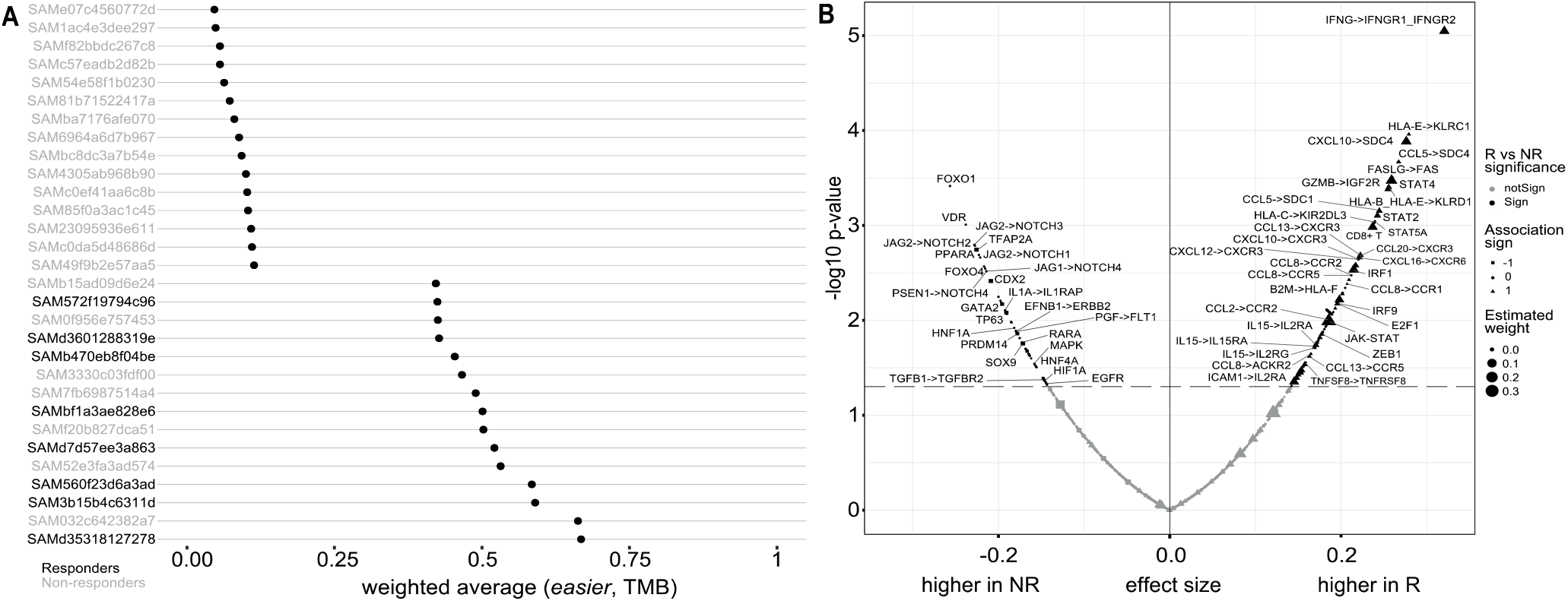
*easier* application. **A.** Patient-specific scores of likelihood of response to ICB therapy based on the weighted average between *easier* predictions and TMB. For visualization, only patients with scores among the top and bottom 15 are shown. Patients’ labels indicate the positive (black) or negative (gray) response to therapy. **B**. Analysis of biomarkers associated with the response to ICB therapy (NR=non-responders, R=responders).

*easier* goes beyond the task of prediction alone, and aims at interpreting predictions through quantitative biomarkers. In fact, this can be of clinical interest, namely to understand why some patients get favorable responses to therapy and others do not. *easier* offers visual solutions to mechanistically interpret patients’ response (Fig. 1B; Supplementary Fig. S4).

## 3 Case study: bladder cancer patients treated with anti-PD-L1 therapy

To illustrate the practical application of *easier*, we re-analyzed a publicly available cohort from patients with metastatic urothelial cancer treated with ICB (Mariathasan et al., 2018). This dataset contains, for 150 patients, bulk RNA-seq pre-therapy data, information on TMB, tumors’ immune phenotype, and observed response to ICB (Complete response=21; Progressive disease =129). In the original study, both pre-existing CD8^+^ T-cell immunity and TMB were suggested as positive indicators of response to ICB, as opposed to the TGFB pathway which plays a negative role in patients’ outcome by contributing to the exclusion of CD8^+^ T cells. Our results showed that combining TMB with *easier* score improved the predictions with respect to the use of TMB as a single predictor (Fig. 1A; Supplementary Fig. S3 and Fig. S5). Consistently, the presence of CD8^+^ T cells was significantly higher in responders’ samples whereas TGFB1→TGFBR2 interaction was significantly over-represented in non-responders (Fig. 1B), especially within tumors from non-responders patients with immune-excluded-phenotypes (Supplementary Fig. S6). Further, *easier* can especially be useful to suggest potential biomarkers that can be followed up experimentally. Several ligand-receptor pairs affecting Notch signaling were enriched in non-responders, confirming the role of Notch in support of an immunosuppressive TME (Colombo et al., 2018). In the context of bladder cancer, a closer look into the crosstalk between Notch and TGFB could depict the mechanisms of T-cell exclusion. Both biomarkers showed higher values in tumors that were both immune-excluded and non-responsive to ICB (Supplementary Fig. S7).

## 4 Conclusions

*easier* is a valuable tool to predict patients’ antitumor immune response that enables mechanistic interpretation of individual patients’ response to treatment based on quantitative biomarkers, which is not possible with standard predictors (e.g. simple gene sets). *easier* leverages widely accessible patients’ tumor bulk RNA-seq data, to characterize the TME from high-level derived features in such manner that resembles information arising from complex and expensive techniques, for instance, imaging for immune-cell quantification or phosphoproteomics upon perturbation for pathway activation. As such, *easier* has potential clinical implications for the scientific community.

## Data and code availability

The data from the bladder cancer cohort (Mariathasan et al., 2018) is made available in IMVigor 210 Biologies R package and can be found at http://research-pub.gene.com/IMvigor210CoreBiologies. *easier* is available under MIT license at https://bioconductor.org/packages/easier as an R/Bioconductor package.

## Supplementary information

**Supplementary Figure S1:**
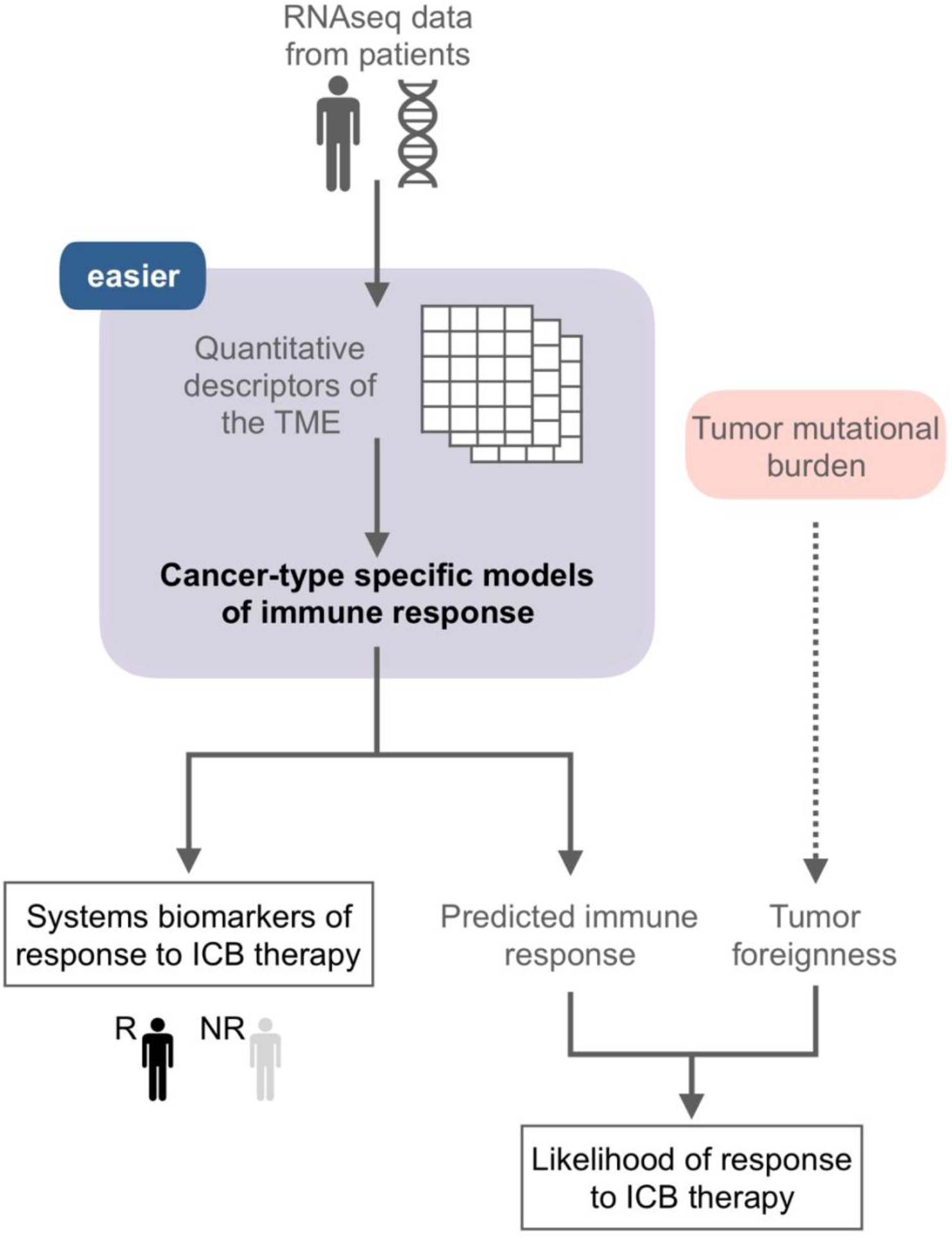
Condensed version of *easier* pipeline.

**Supplementary Figure S2:**
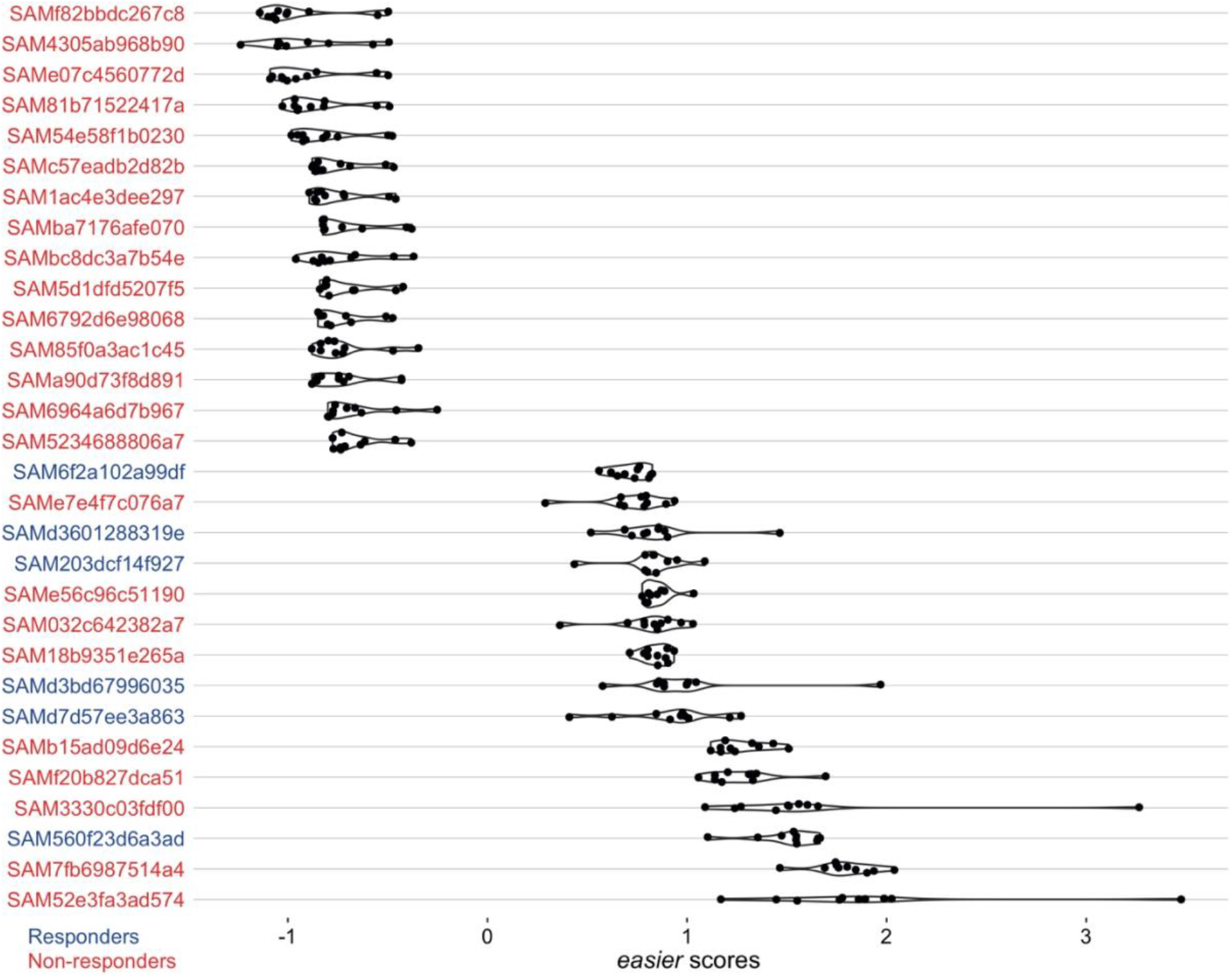
*easier* scores for each patient. Each dot corresponds to each task (i.e. different hallmarks of immune response). Patients are colored according to their positive (blue) or negative (red) response to anti-PD-L1 therapy. Patients are sorted according to the median across tasks. For visualization purposes, only patients with median values among the top and bottom 15 are shown.

**Supplementary Figure S3:**
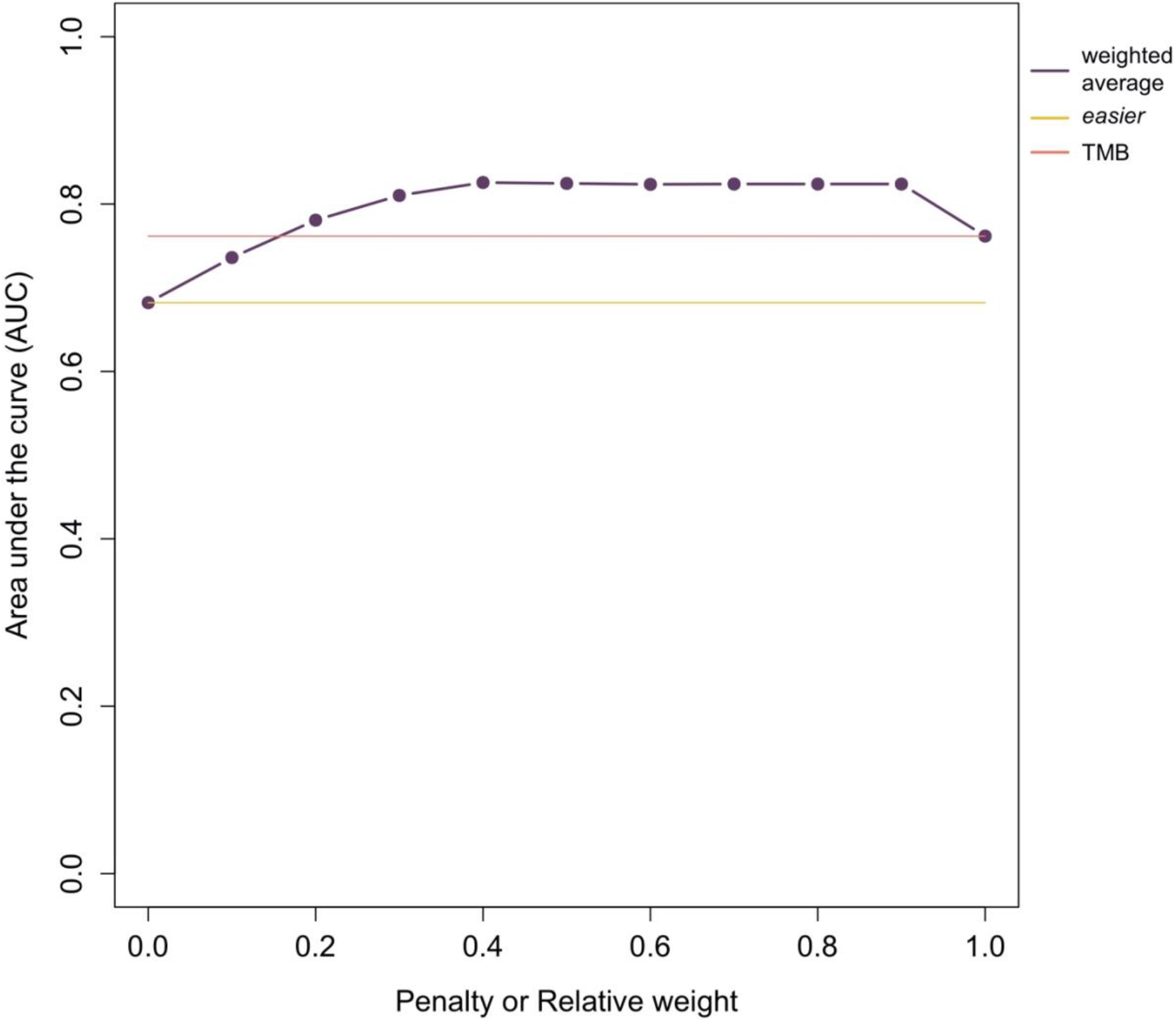
Comparison of the performance of the predictions, calculated using the Area under the curve, for *easier*, TMB and the integrated score (i.e. weighted average). The integrated score is based on the weighted average between *easier* and TMB score. The integrated score is shown for different weight values.

**Supplementary Figure S4:**
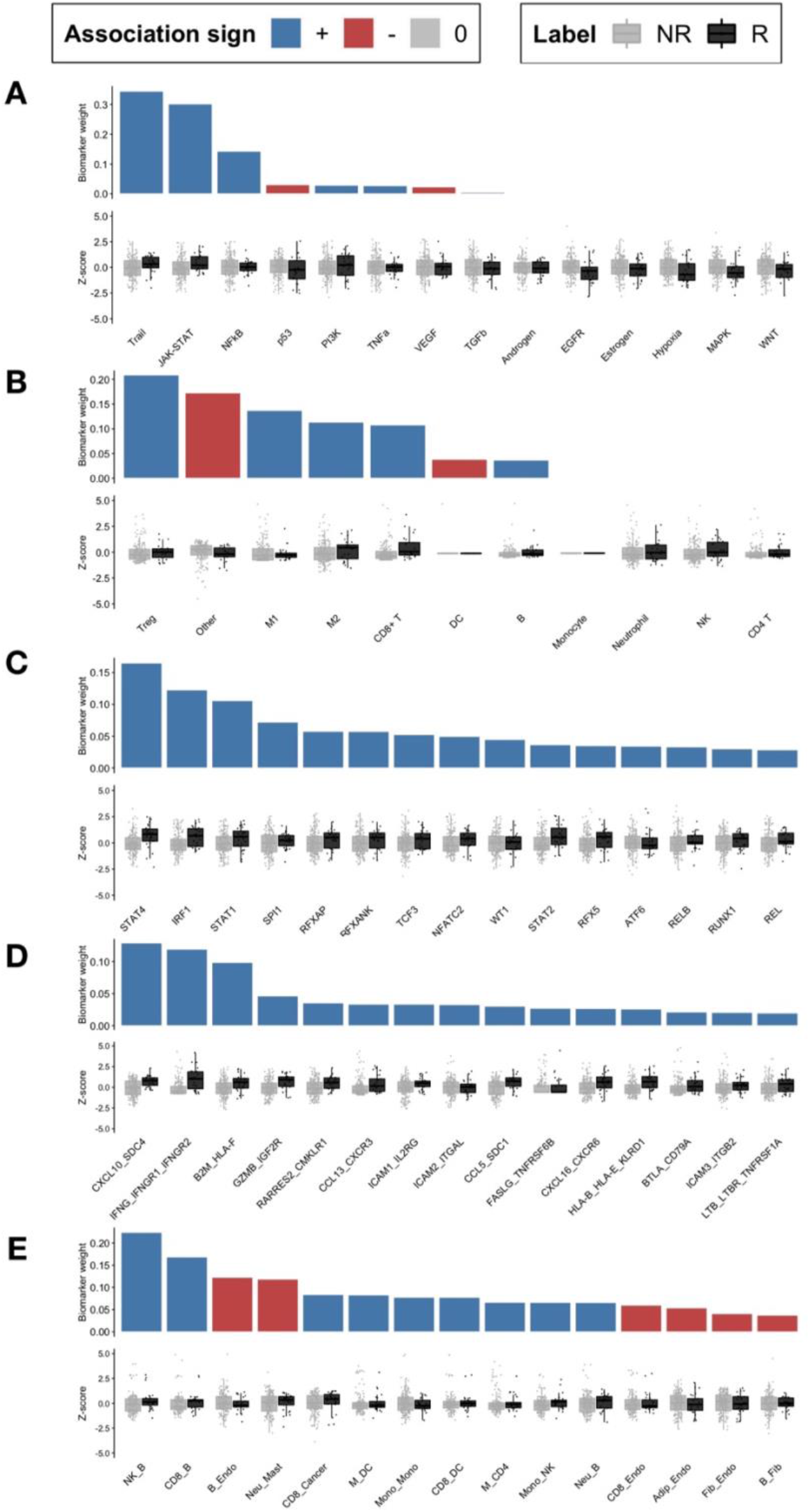
Quantitative biomarkers based on **A.** Pathway activity, **B.** Immune cell quantification, **C.** Transcription Factor activity, **D.** Ligand-Receptor pairs and **E.** Cell-Cell scores. Box plots displayed biomarker values comparing non-responder (NR) and responder patients (R). Bar plots showed the regression coefficient for each biomarker. Biomarkers are sorted according to their absolute regression coefficient value.

**Supplementary Figure S5:**
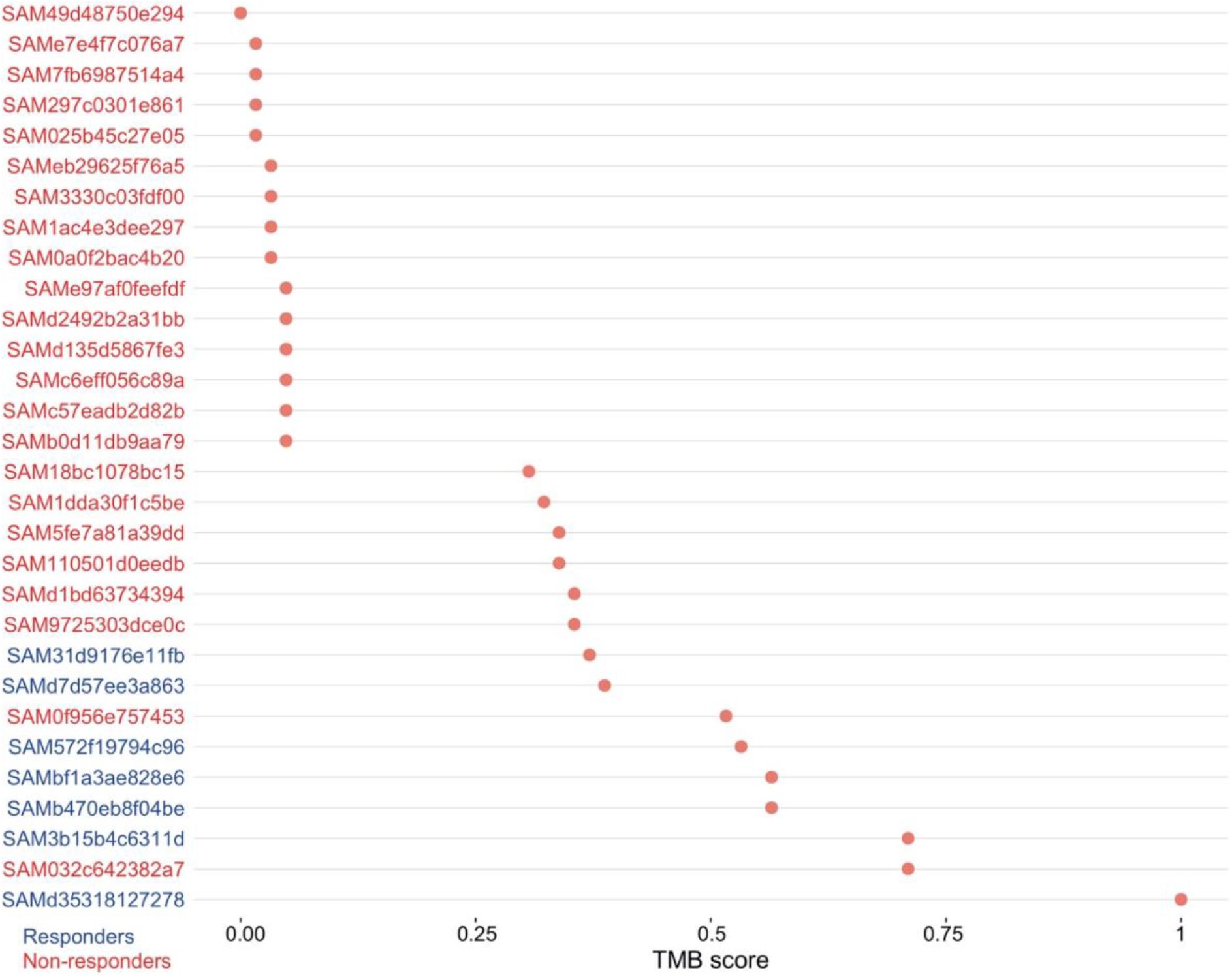
Tumor mutational burden (TMB) scores for each patient. Patients are sorted according their TMB score. For visualization purposes, only patients with TMB score among the top and bottom 15 are shown. Patients are colored according to their positive (blue) or negative (red) response to anti-PD-L1 therapy.

**Supplementary Figure S6:**
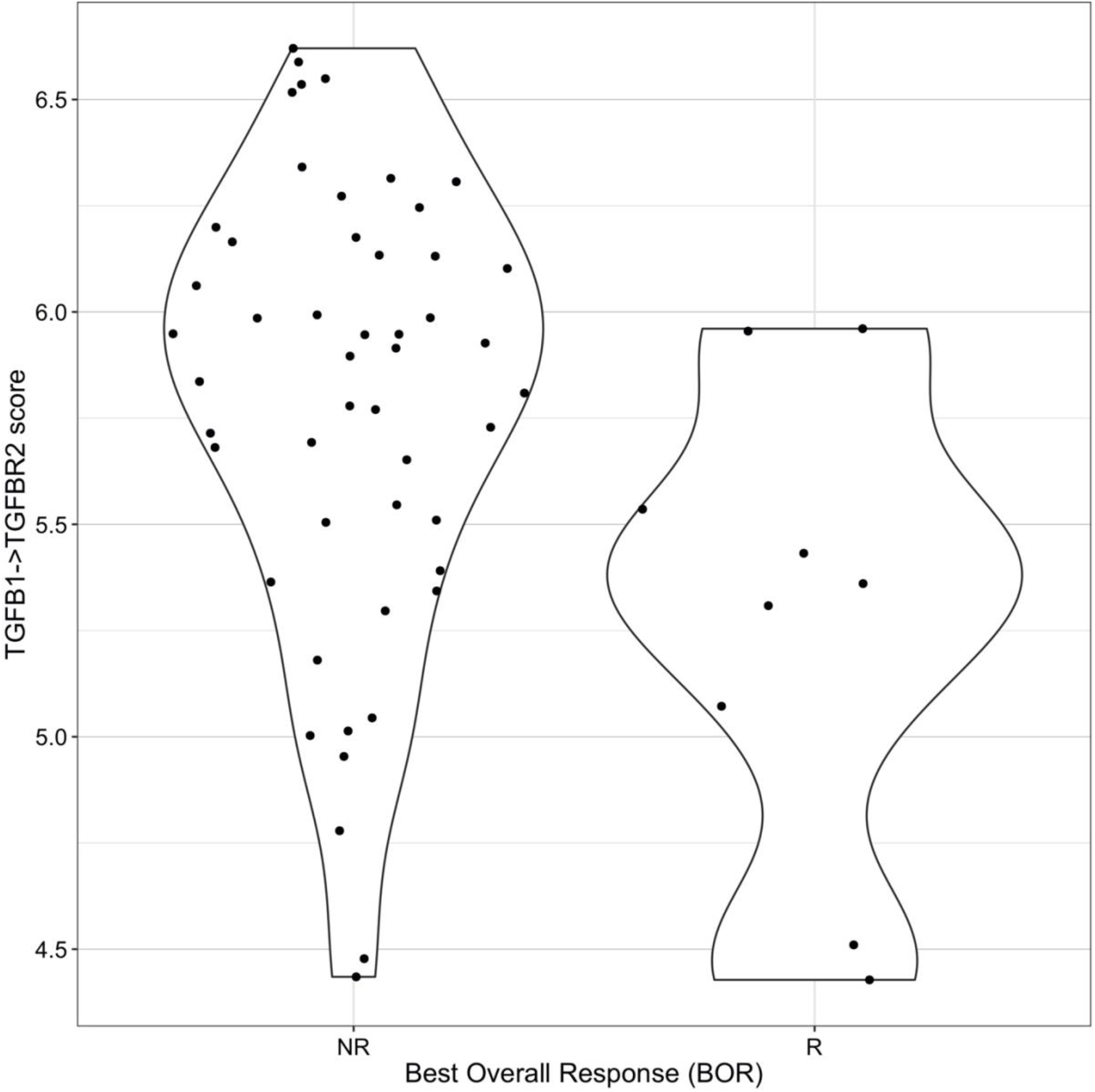
Comparison of TGB1→TGFB2 ligand-receptor pair feature values across non-responders (NR) and responders (R) from immune-excluded tumors.

**Supplementary Figure S7:**
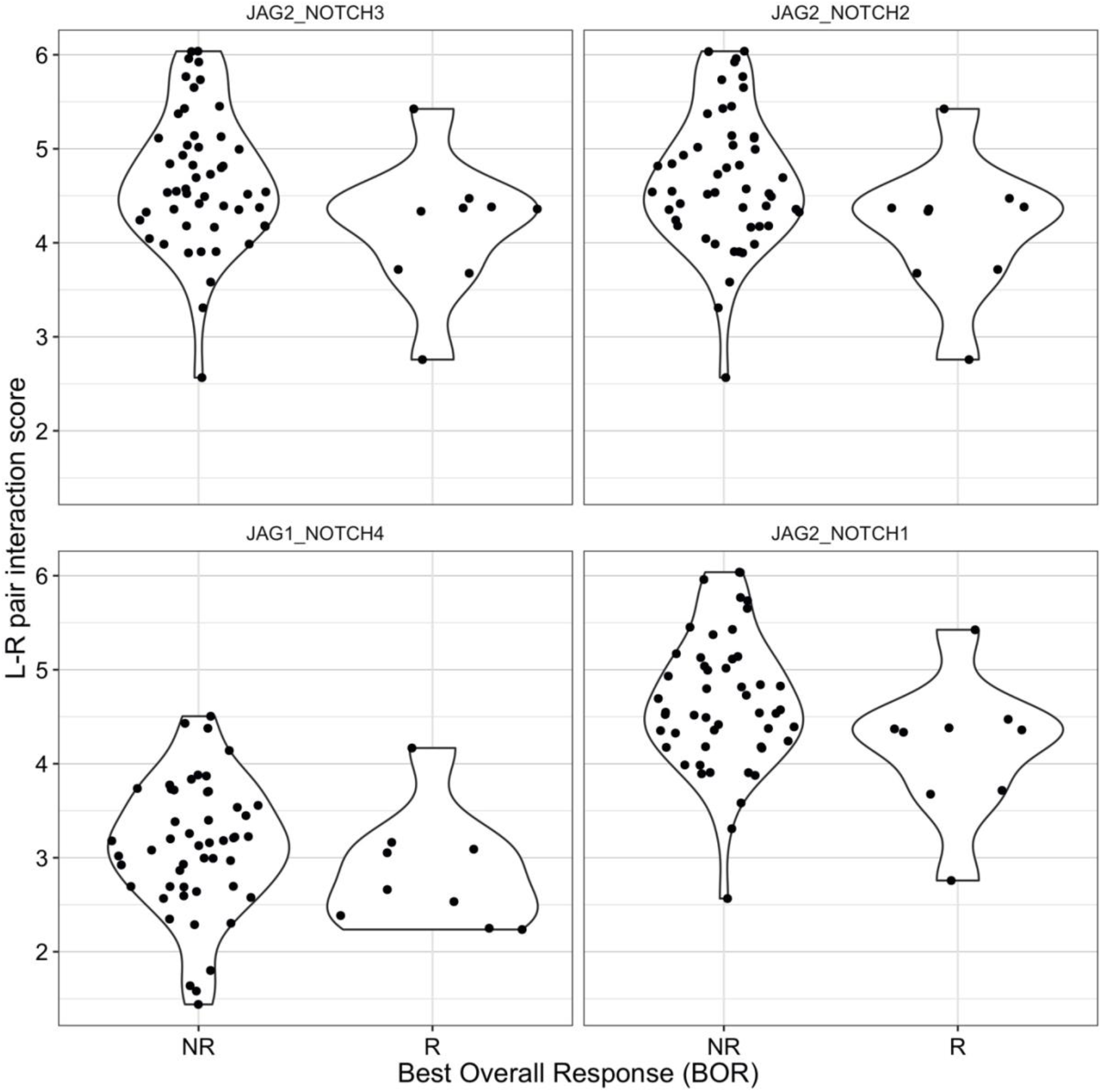
Comparison of JAG1→NOTCH4, JAG2→NOTCH3, JAG2→NOTCH2 and JAG2→NOTCH1 ligand-receptor pair features values across non-responders (NR) and responders (R) from immune-excluded tumors.

